# Genome assembly of *Hibiscus sabdariffa* L. provides insights into biosynthetic pathways of medicinal natural products

**DOI:** 10.1101/2023.07.10.548471

**Authors:** Taein Kim, Jeong Hun Lee, Hyo Hyun Seo, Sang Hyun Moh, Sung Soo Choi, Jun Kim, Sang-Gyu Kim

## Abstract

*Hibiscus sabdariffa* L. is a widely cultivated herbaceous plant with diverse applications in food, tea, fiber, and medicine. In this study, we present a high-quality genome assembly of *H. sabdariffa* using more than 33 Gbps of high-fidelity (HiFi) long-read sequencing data at 20X coverage. The primary assembly and haplotype-resolved genomes exhibit an N50 contig length of 26.25 Mbps, 11.96 Mbps, and 14.50 Mbps, with a coverage of 141.3%, 86.0%, and 88.6%, respectively. By aligning 26 Gbps of total RNA sequencing data (PE100) extracted from leaves, we predicted 154k, 79k, and 87k genes in the respective genome. The completeness of the genomes and predicted genes was confirmed by benchmarking universal single-copy orthologs analysis on a eudicots_odb10 database, with a 99.3% complete rate for the primary assembly. The high-quality genome resource allowed the construction of genetic networks for phenylpropanoid and flavonoid metabolism, thereby facilitating the identification of candidate genes involved in the biosynthesis of key intermediates for roselle-specific medicinal natural products. Comprehensive genomic and functional analysis opens avenues for further exploration and application of valuable natural products derived from *H. sabdariffa*.

## Introduction

*Hibiscus*, a popular genus in the Malvaceae family, contains more than 200 species distributed in tropical and subtropical regions. Plants in genus *Hibiscus* are widely known to produce valuable natural products such as fibers, pigments, and medicinal secondary metabolites. The ploidy levels of the genus *Hibiscus* vary, with examples such as *Hibiscus rosa-sinensis* having 2n = 16x = 144 and *Hibiscus syriacus* having 2n = 4x = 80 (Zhang *et al*. 2020). *Hibiscus sabdariffa* or “roselle” having a ploidy of 2n = 4x = 72 is widely cultivated in tropical climates such as Sudan, India, South America, and China (Sarhadynejad *et al*. 2013). High-quality genomes of *Hibiscus mutabilis*, *H. syriacus*, and *Hibiscus trionum* have been published (Li *et al*. 2015; Kim *et al*. 2017; Yang *et al*. 2022; Koshimizu *et al*. 2023). In contrast, the available genome for *H. sabdariffa* is currently limited to a poor-quality draft, which poses a challenge for comprehensive genomic and biochemical studies in roselle. (Kim *et al*. 2017; Yang *et al*. 2022; Koshimizu *et al*. 2023).

As a medicinal plant, *H. sabdariffa*’s phytochemical and pharmacological properties are well studied (Da-Costa-Rocha *et al*. 2014). The calyx of *H. sabdariffa* contains various bioactive organic acids, including phenolic acid, hydroxycitric acid, and hibiscus acid. In addition, *H. sabdariffa* contains anthocyanins and flavonoids such as delphinidin, cyanidin, hibiscetin, gossypetin, and quercetin. (Da-Costa-Rocha *et al*. 2014). These bioactive compounds have been the subject of considerable research interest due to their potential health benefits and therapeutic properties. In *H. sabdariffa* extract, quercetin and its derivative, quercetin-3-O-glucuronide, have been found to decrease the levels of reactive oxygen species (ROS) induced by glucolipotoxicity. They also reduce triglyceride accumulation, which is critical for anti-obesity (Herranz-López *et al*. 2020). In addition, the roselle extracts showed antihypertensive effect, anti-inflammatory effect, and antimicrobial effect (Herrera-Arellano *et al*. 2004; Mardiah *et al*. 2015; Abdallah 2016).

Revealing the biosynthetic pathways of key natural products responsible for these medicinal effects would facilitate the pharmacological application of *H. sabdariffa*. Recently, transcriptomics-based approaches have been developed as effective tools for revealing the synthetic mechanism of complex natural products (Nett *et al*. 2020; Kang *et al*. 2022; Sun *et al*. 2022b; Li *et al*. 2023; Alagna *et al*. 2023). The availability of a high-quality genome resource of the host organism is essential for reliable transcriptome analysis and manipulation of gene expression (Raymond *et al*. 2018; Saint-Oyant *et al*. 2018; Sun *et al*. 2022b; Li *et al*. 2023). Moreover, comparative genomics studies provide insights into evolutionary aspects and functional diversification, providing critical evidence for the biogenesis of novel natural products (Zhang *et al*. 2020, 2023; Sun *et al*. 2022a; Tang *et al*. 2022; Wang *et al*. 2023; Conart *et al*. 2023).

In this study, we reported a high-quality *H. sabdariffa* genome and pseudo-haplotype genome, constructed using the high-fidelity (HiFi) long-read sequencing technology of Pacific Biosciences (PacBio). The quality of genomes and annotated genes was validated by length distribution and BUSCOs analysis (Fig. 1 & Table 1). Functional analysis was conducted to identify genes involved in natural product biosynthetic pathways. The availability of these resources and the implementation of functional analysis present opportunities for biochemical understanding of *H. sabdariffa* and its natural products.

**Figure 1.**
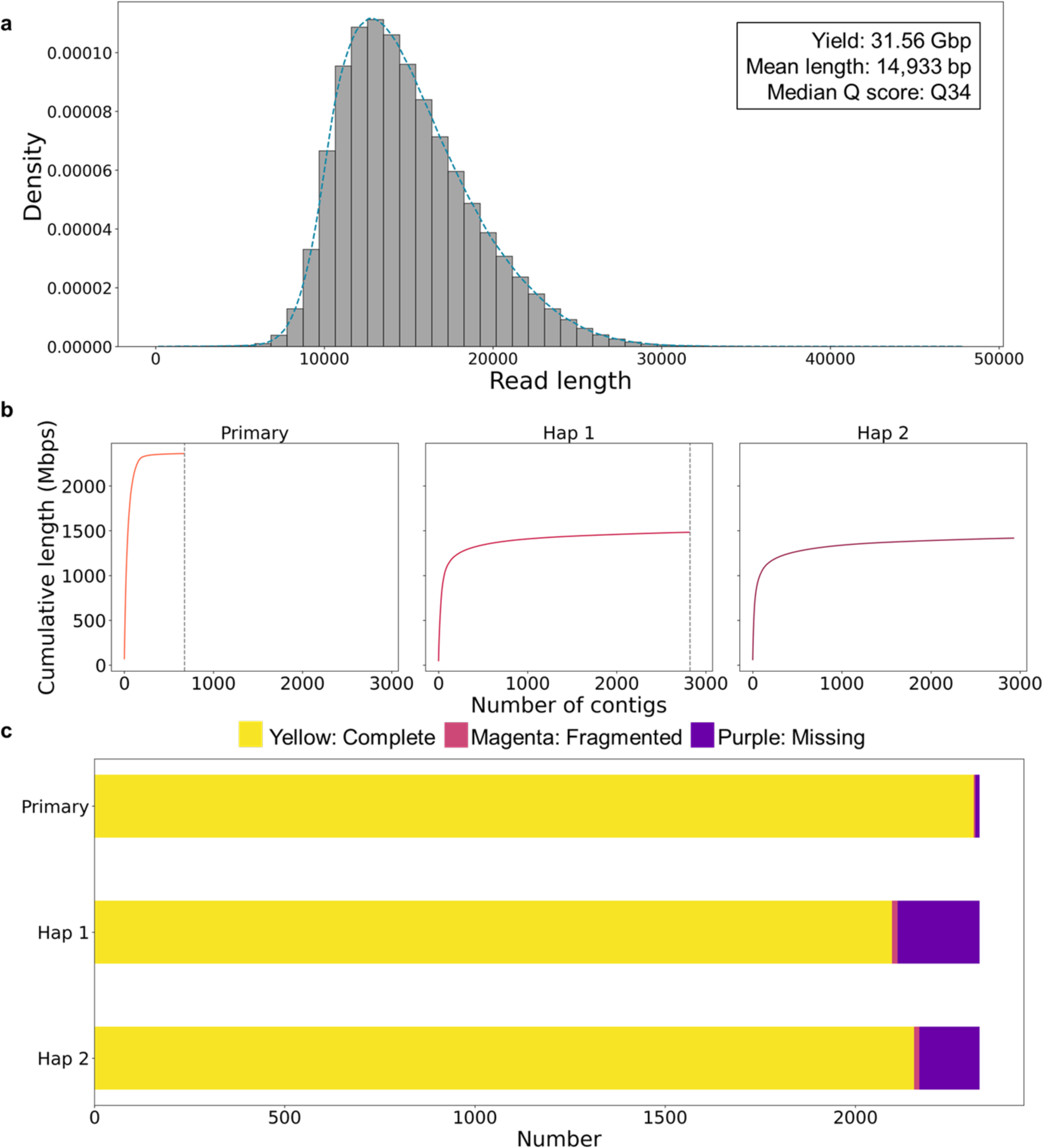
a) The read length distribution of *Hibiscus sabdariffa* genomic DNA sequenced by PacBio HiFi sequencing. b) The cumulative length of contigs of haplotypes in descending order. Grey lines represent the numbers of contigs in primary assembly and Hap 1. c) The BUSCO scores of primary assembly, Hap 1, and Hap 2 genome using eudicots_odb10 database.

**Table 1.** The raw sequencing data and assembled genomes of *Hibiscus sabdariffa*

| Long-read genomic DNA sequencing |  |  |  |  |
| --- | --- | --- | --- | --- |
| Type | Read Count | Yield (bp) | Mean Length (bp) | Mean Q score |
| PacBio HiFi sequencing | 2,269,272 | 33,887,858,605 | 14,993 | Q34 |
| RNA sequencing |  |  |  |  |
| Type | Read Count | Yield (bp) | Mean Length (bp) | > Q30 |
| Illumina PE100 | 263,077,312 | 26,570,808,512 | 100 | 94.8 % |
| Assembled genomes of *Hibiscus sabdariffa* |  |  |  |  |
|  | Primary | Haplotig 1 | Hapltig 2 |  |
| Number of Contigs | 675 | 2929 | 2823 |  |
| Max Contig (Mbp) | 70.47 | 60.86 | 48.4 |  |
| Mean Contig (Mbp) | 3.50 | 0.48 | 0.53 |  |
| Contig N50 (Mbp) | 26.25 | 11.96 | 14.5 |  |
| Total Length (Gbp) | 2.36 | 1.42 | 1.48 |  |

## Materials and methods

### Plant Materials

Commercially available *Hibiscus sabdariffa* seeds (Cheonnyangssiat, Chungcheongbuk-do, South Korea) originated from Hongkong were used for genome assembly and annotation. The seeds were germinated and were grown in plant growth room of 16 h-8 h light-dark photoperiod with controlled temperature at 25 ± 2 °C and moiety at 60 %. The adult leaves of the plants were sampled and delivered to NICEM (Seoul National University, South Korea) for DNA sequencing. The adult leaves were collected for RNA sequencing.

#### DNA/RNA sequencing

The genomic DNA of *H. sabdariffa* plants was extracted from adult leaves using a following protocol. The extraction method employed was based on the cetyltrimethylammonium bromide (CTAB) method, known for its high molecular weight DNA extraction capabilities. The extraction procedure involved sample preprocessing, cell lysis, DNA purification, DNA concentration, and sequential elimination of short DNA to ensure the quality and integrity of the extracted genomic DNA. Extracted genomic DNA was qualified with Nanodrop, Quantus, and agarose gel electrophoresis to validate whether its quality is adequate to be sequenced. The PacBio HiFi sequencing libraries were prepared with SMRTbell prep kit 3.0 (Pacific Biosciences, Menlo Park, USA) and were sequenced on the SMRT bell barcoded adapter plate (Pacific Biosciences) and PacBio Sequel IIe (Pacific Bioscience) at NICEM.

Total RNAs of *H. sabdariffa* adult leaves were extracted with the RNeasy Plant Mini Kit (Qiagen, Hilden, Germany), and RNase-free DNase I (Qiagen) was used to eliminate the genomic DNA from extracted RNA. Quality of RNA was checked with RNA integrity number score on 2100 Bioanalyzer (Aligent Technologies, Santa Clara, USA). The RNA sequencing libraries were generated with the TruSeq Stranded Total RNA with Ribo-Zero Plant kit (Illumina, San Diego, USA) and were sequenced on NovaSeq (Illumina) at Macrogen (Seoul, South Korea).

#### De novo genome assembly

Genome assembly and annotation were performed following a protocol paper (Kim and Kim 2022). The HiFi sequencing reads of *H. sabdariffa* genomic DNA were *de novo* assembled into contig level genome using the Hifiasm assembler (v 0.18.5-r500, cloned from official GitHub repository in February 14, 2023) on defaults options except -s 0.54 (-s is the option indicating threshold of duplicate haplotigs in real-level), and the haplotigs (Hap1 and Hap2 in this paper) of the draft genome were separated in assembly procedure of Hifiasm (Cheng *et al*. 2021). The graphical fragment assembly format (GFA) files contained in Hifiasm results were converted into FASTA format using the “awk” command in Linux. This conversion process involved extracting the sequence information from the GFA files and organizing it into the standard FASTA format. The genomic sequence of each haplotig was treated individually for all following analysis.

#### Repeat and gene annotation

Since there is no available specific repeat library for *H. sabdariffa*, we constructed *de novo* repeat library of *H. sabdariffa* using RepeatModeler (v2.0.3) with NINJA (NINJA-0.95-cluster_only)(Jullien *et al*. 2020). RepeatMasker (v4.1.2) masked repetitive regions of primary assembly and both haplotigs based on pre-defined eudicots repeat library and the *de novo* repeat library with defaults options(Saha *et al*. 2008). This two-step approach ensures comprehensive masking of repetitive elements in the genomic sequences, enhancing the accuracy of downstream analyses. The results of hard masking were used for analysis of repetitive regions and the result files of soft masking were further utilized to gene annotation.

Based on repeat masking results, we found the telomeric repeats in the genome applying “grep” command to find the plant canonical telomeric repeat sequence “TTTAGGG” and its variant forms (Richards and Ausubel 1988; Adamusová *et al*. 2020). The candidate telomeric regions were inferred by the position and length, i.e., the telomeric repeats placing in the end of contig and longer than 1000 bps (i.e. ∼140 concatamers of telomeric repeat) were considered as candidate telomeres. When the position of telomeric repeats is 5’ end, the sequence should be “CCCTAAA” and its variant forms, and “TTTAGGG” and its variant forms were required for 3’ end.

Total RNA sequencing reads were aligned to the soft-masked genomes with HISAT2 (v2.2.1) and the alignment results were sorted into the binary alignment and map (BAM) formats with SAMtools (v1.15 using htslib 1.14) (Li *et al*. 2009; Kim *et al*. 2019). Based on the RNA-seq alignment results and *ab-initio* gene prediction model, genes were annotated for each soft masked genome with BRAKER2 (v2.1.6 using Genemark-ES/ET/EP v.4.71_lic) default options (Stanke *et al*. 2006, 2008; Li *et al*. 2009; Barnett *et al*. 2011; Buchfink *et al*. 2014; Lomsadze *et al*. 2014; Hoff *et al*. 2016; Brůna *et al*. 2020, 2021).

#### Evaluation

Qualities of assembled genome and annotated genes were evaluated based on statistics and benchmarking universal single-copy orthologs (BUSCO) results (Simão *et al*. 2015). BUSCO (v.5.2.2) analysis was performed on the eudicots_odb10 database to confirm whether assembled genomes and predicted gene sets appropriately contain the evolutionary conserved genes (Simão *et al*. 2015). We employed the assembly-stats (v1.0.1) to analyze the length statistics of contigs and genes, including the number, mean length, maximum length, N50 value, and total size.

#### Structural variants and single nucleotide polymorphism calling

We examined structural variants (SVs) between haplotigs and single nucleotide polymorphisms (SNPs) among primary assembly and haplotigs. Since the Hap 2 showed the longer N50 length than Hap 1, Hap 2 was treated as reference genome in SV calling. Minimap2 (v2.24-r1122) was used to map the Hap 1 assembly to Hap 2, followed by sorting to generate a genome mapping BAM file using SAMtools (Li *et al*. 2009; Li 2018). Based on the genome-to-genome mapping BAM file, svim-asm (v1.0.3) called the SVs between two haplotigs, generating variant call format (VCF) files (Heller and Vingron 2020). By aligning these VCF files to protein coding regions of the genomes using BEDTools (v2.27.1), genes related with SVs were selected (Quinlan and Hall 2010). To call SNPs, HiFi sequencing long-reads of genomic DNA previous used in *de novo* genome assembly were mapped to each primary assembly and haplotigs using minimap2 and SAMtools (v2.24-r1122 and v1.15, respectively) (Li *et al*. 2009; Li 2018). DeepVariant-gpu (v1.5.0) was utilized as a SNP caller in primary assembly and each haplotype based on the mapping files, producing the VCF result files (Poplin *et al*. 2018). The VCF files were annotated with SnpEff (v5.1d) to predict the effect of SNPs (Cingolani *et al*. 2012) on genes.

#### Genetic metabolic network construction

The coding sequences (CDS) of genes predicted in primary and haplotypes genome were functionally annotated by similarity-based sequence search on UniProt Knowledgebase (release 2023.01) using MMseqs2 (v13.4511) (Bateman *et al*. 2017; Steinegger and Söding 2017). The major metabolic profile of *H. sabdariffa* was defined based on PubChem taxonomy data (Supplementary Data 1), the referenced compound-organism pair database (Kim *et al*. 2021). To construct genetic metabolic network of the major metabolic profile of roselle, we referred the corresponding molecular pathways of the KEGG database (Kanehisa and Goto 2000). Subsequently, we extracted annotations of biosynthetic enzymes involved in the production of key intermediate compounds from pathways in other organisms using the KEGG database (Kanehisa and Goto 2000). To identify potential candidate genes for biosynthetic pathways in *H. sabdariffa*, we manually compared these annotations of interests to the functional gene annotations of *H. sabdariffa* genes, constructing a gene set potentially involved in the phenylpropanoids and flavonoids pathways. This approach allowed us to narrow down and prioritize the candidate genes for further investigation and characterization in the context of biosynthetic pathways in *H. sabdariffa*.

## Results and discussion

### Genome assembly and annotation

PacBio HiFi sequencing of *H. sabdariffa* genomic DNA resulted in 2.3 million reads (total 33.9 Gbps); those mean length was 15.0 Kbps (Table 1). The HiFi sequencing read quality of *H. sabdariffa* genomic DNA was verified by a Phred quality score (Q score) of Q34. This high-quality score assures the reliability and confidence of the sequencing data, allowing accurate downstream analyses and interpretations of the genomic information of *H. sabdariffa*. The karyotype of *H. sabdariffa* is 2n = 4x = 72, and the haplotype genome size is 1.67 Gbps, i.e., we got 20.3X coverage raw reads (Mohammad *et al*. 2020). The HiFi reads were assembled with Hifiasm and phased into pseudo haploids, Hap 1 and Hap 2 (Cheng *et al*. 2021). The primary assembly represented a total length of 2.36 Gbps with a contig N50 length of 26.25 Mbps and the longest contig measuring 70.47 Mbps (Table 1). For the resolved haplotigs, the total length, N50 length, and maximum length of contigs were found to be 1.42 Gbps, 11.96 Mbps, and 60.86 Mbps for Hap1, and 1.48 Gbps, 14.50 Mbps, and 48.40 Mbps for Hap2, respectively (Table 1). The coverage of the primary genome, Hap1 genome, and Hap2 genome were 141.3%, 86.0%, and 88.6%. In this study, Hap2 was used as a reference haplotype of *H. sabdariffa* genome for structural variation analysis.

We evaluated each genome with respect to the length distribution of contigs and BUSCOs score(Simão *et al*. 2015). The cumulative contig length graphs of the assembled genomes exhibited a left-skewed shape (Fig. 1), indicating a higher proportion of longer contig lengths. The contig N50 length of 26.25 Mbps for primary genome (consisted of 28 contigs) was comparable to 46.39 Mbps, the arithmetic average of chromosome lengths (A total genome size / the number of chromosomes) of *H. sabdariffa*. The completeness of genome was further evaluated by BUSCO analysis on eudicots_odb10 database, which represented 99.3% complete rate for primary genome, 90.1% for Hap 1, and 92.6% for Hap 2 (Fig. 1c), which are comparable with the chromosome-level genomes of other species in *Hibiscus* genus (Simão *et al*. 2015; Yang *et al*. 2022).

The tetraploid nature of *H. sabdariffa* poses a challenge in *de novo* genome assembly, occasionally leading to mis-assemblies. Hifiasm is renowned for its ability to preserve the contiguity of haplotypes during genome assembly as a leading haplotype-resolved assembler (Cheng *et al*. 2021). In this study, we performed both primary assembly and haplotype-resolved assembly. The primary assembly could represent each homologous sequence once and each heterologous “bubble” twice. The primary genome size of *H. sabdariffa* was 2.36 Gbps, which is larger than the haplotype genome (1.67 Gbps) but smaller than the diploid genome (3.34 Gbps) of *H. sabdariffa*. However, the lower BUSCO score observed in the haplotypes compared to the primary assembly might be due to the inherent challenges in resolving a complexity in tetraploid genome. On the other hand, the genome sizes of resolved haplotigs (Hap 1 and Hap 2) were smaller than the theoretical haplotype genome size (1.42 and 1.48 Gbps, respectively). This difference can be explained by the fact that each haplotype is a reduced version of the primary assembly of the 2X genome (n = 2x = 36). To enhance the phasing and contiguity of the *H. sabdariffa* genome, incorporating parental sequencing information and Hi-C data would improve the resolution of haplotypes, leading to a more comprehensive and accurate representation of the genome. It is worthy to mention that despite this limitation, Hifiasm is a viable solution due to its graph-binning strategy and its capacity for constructing haplotype-resolved assemblies (Cheng *et al*. 2021; Sun *et al*. 2022a).

We next performed repeat masking with the eudicot repeat library of ReapeatMasker. Lower than 4% of genome were annotated as simple and low complexity repeats. On the other hand, after being masked with *de novo* modeled repeat library, a significant portion of genome was annotated as repeats: 74.48% in primary genome, 76.67% in Hap 1, and 73.99% in Hap 2 (Fig. 2a, Tables 2 & 3). Hap 2 genome consists of 42.91% retroelements, 1.10% DNA transposons and 24.43% unclassified repeats, where Hap 1 consists of 46.56% retroelements, 0.97% DNA transposons, and 23.05% unclassified repeats (Table 3). Specifically, more than 40% regions (41.69% in primary assembly, 44.59% in Hap 1, and 41.42% in Hap 2) of genomes were LTR elements, containing Ty1/Copia, Gypsy/DIRS1, and retroviral sequences (Table 3).

**Figure 2.**
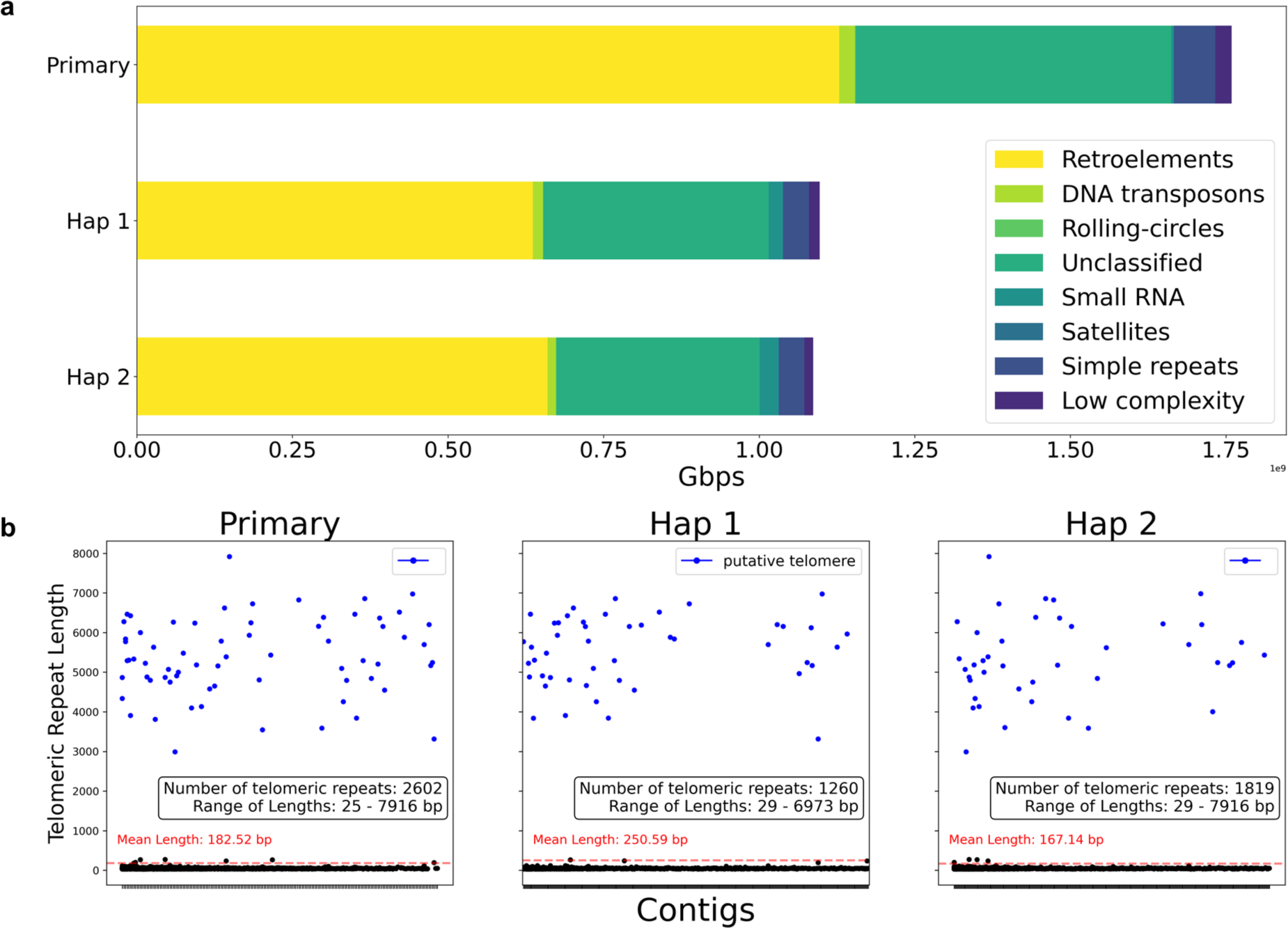
a) The composition of repeat components in *H. sabdariffa* genome. b) The telomeric repeats distribution in *H. sabdariffa* genome.

**Table 2.** Repetitive element annotations of *H. sabdariffa* primary assembly

| Repetitive elements of primary assembly |  |  |  |
| --- | --- | --- | --- |
| Type | Number of elements | Length (bp) | Coverage in genome |
| Retroelements | 923,542 | 1,128,617,308 | 47.79% |
| DNA transposons | 38,811 | 25,253,436 | 1.07% |
| Rolling-circles | 1,725 | 971,658 | 0.04% |
| Unclassified | 1,678,392 | 507,172,818 | 21.47% |
| Small RNA | 13,660 | 4,066,571 | 0.17% |
| Satellites | 539 | 184,424 | 0.01% |
| Simple repeats | 1,014,176 | 656,886,412 | 2.83% |
| Low complexity | 289,769 | 26,090,467 | 1.1% |
| Total | 3,960,614 | 1,759,043,094 | 74.48% |
| Retroelements |  |  |  |
| Type | Number of elements | Length (bp) | Coverage in genome |
| SINES | 932 | 51058 | 0.00% |
| penelope | 3382 | 2160750 | 0.09% |
| LINEs | 74279 | 84394825 | 3.57% |
| LTR elements-Ty1/Copia | 424379 | 606739301 | 25.69% |
| LTR elements-Gypsy/DIRS1 | 263913 | 377767974 | 16.00% |
| LTR elements-Retroviral | 82 | 20714 | 0.00% |

**Table 3.** Repetitive element annotations of *H. sabdariffa* haplotype-resolved genome

| Repetitive elements of Hap 1 |  |  |  |
| --- | --- | --- | --- |
| Type | Number of elements | Length (bp) | Coverage in genome |
| Retroelements | 458,524 | 659,844,473 | 46.56 % |
| DNA transposons | 22,207 | 13,703,639 | 0.97 % |
| Rolling-circles | 1,504 | 458,855 | 0.03 % |
| Unclassified | 1,203,054 | 326,717,778 | 23.05 % |
| Small RNA | 13,868 | 30,709,060 | 2.17 % |
| Satellites | 128 | 47,021 | 0.00 % |
| Simple repeats | 603,428 | 41,007,044 | 2.89 % |
| Low complexity | 163,171 | 14,169,642 | 1.00 % |
| **Total** | **2,465,884** | **1,086,657,512** | **76.67%** |
| Retroelements |  |  |  |
| Type | Number of elements | Length (bp) | Coverage in genome |
| SINES | 13,160 | 1,678,841 | 0.12 % |
| LINEs | 36,529 | 26,299,893 | 1.86 % |
| LTR elements-Ty1/Copia | 187,929 | 320,865,058 | 22.64 % |
| LTR elements-Gypsy/DIRS1 | 160,992 | 259,764,164 | 18.33 % |
| LTR elements-Retroviral | 25,920 | 40,339,228 | 2.85 % |
| LTR elements-Others | 33,994 | 10,897,289 | 0.77 % |
| Repetitive elements of Hap 2 |  |  |  |
| Type | Number of elements | Length (bp) | Coverage in genome |
| Retroelements | 464,120 | 636,317,063 | 42.91 % |
| DNA transposons | 26,861 | 16,281,690 | 1.10 % |
| Rolling-circles | 1,781 | 473,989 | 0.03 % |
| Unclassified | 1,414,803 | 362,236,882 | 24.43 % |
| Small RNA | 12,182 | 22,910,841 | 1.54 % |
| Satellites | 104 | 41,919 | 0.00 % |
| Simple repeats | 637,863 | 41,654,744 | 2.81 % |
| Low complexity | 188,519 | 17,334,693 | 1.17 % |
| **Total** | **2,746,233** | **1,097,251,821** | **73.99%** |
| Retroelements |  |  |  |
| Type | Number of elements | Length (bp) | Coverage in genome |
| SINES | 15,624 | 1,996,449 | 0.13 % |
| LINEs | 42,506 | 29,505,767 | 1.99 % |
| LTR elements-Ty1/Copia | 184,357 | 287,549,490 | 19.39 % |
| LTR elements-Gypsy/DIRS1 | 165,733 | 280,261,487 | 18.9 % |
| LTR elements-Retroviral | 17,479 | 24,749,986 | 1.67 % |
| LTR elements-Others | 38,391 | 12,253,884 | 0.83 % |

The telomeric repeats of *H. sabdariffa* were analyzed using repeat masking results. The telomeric repeat sequence of *H. sabdariffa* genome was “TTTAGGG”, which is known as the canonical repeat sequence of plants (Richards and Ausubel 1988). *H. sabdariffa* has 72 telomeric regions in its chromosome. Among the 2602 telomeric repeat regions in primary assembly, 67 repeats were inferred to putative telomere that were positioned at the end of each contigs and longer than 1,000 bps (∼140 concatamers) (Fig. 2b). In Hap 1 and Hap 2, 47 and 42 repeats were annotated as a putative telomere (Fig. 2b). The longest putative telomeres were 7,916bp in primary assembly, 6,973 bps in Hap 1, and 7,916 bps in Hap 2 (Fig. 2b).

We produced about 26 Gbps of total RNA sequencing reads, which of 94.8% were higher than Q30 quality (Table 1). Each soft-masked genome was gene-annotated hinted by aligning those total RNA sequencing data onto each genome. 154,226 genes in primary assembly, 79,143 genes in Hap 1, and 87,414 genes in Hap 2 were annotated (Table 1). The mean and N90 length of genes were 1,033 bps and 480 bps for primary assembly, 1,074 bps and 510 bps for Hap 1, and 1,099 bps and 525 bps for Hap 2, implying the validity of the annotated genes (Table 3). Each annotated genes set was translated into protein sequence and validated by protein BUSCO with the same database to genome BUSCO (eudicots_odb10 database). The completes rates were 99.3% for primary assemble genome, 88.9% for Hap 1, and 92.0% for Hap 2 (Table 3).

### SNP (single nucleotide polymorphism) and SV (structural variation) calling

SNPs in primary assembly, Hap 1, and Hap 2 genomes were analyzed for measuring genomic variation. The impact of SNP on protein structure was classified into three categories: high, moderate, and low following the criteria of SnpEff (Cingolani *et al*. 2012). In the primary assembly, Hap 1, and Hap 2 genomes, we found 496, 6,673, and 6,147 of high impact SNPs, 10,196, 330,362, and 301,055 of moderate impact SNPs, and 6,147, 439,609, and 399,528 of low impact of SNPs, respectively (Fig. 3b). Subsequently, the SVs between Hap 1 and Hap 2 genomes were called and intersected with gene annotation to find out gene sets that were affected by these SVs. Hap 2 genome contained 341, 194, and 75 genes related with deletion, insertion, and PAVs, respectively (Fig. 3a). Given the redundancy observed in tetraploid organisms, these genes are expected to have the potential for neofunctionalization.

**Figure 3.**
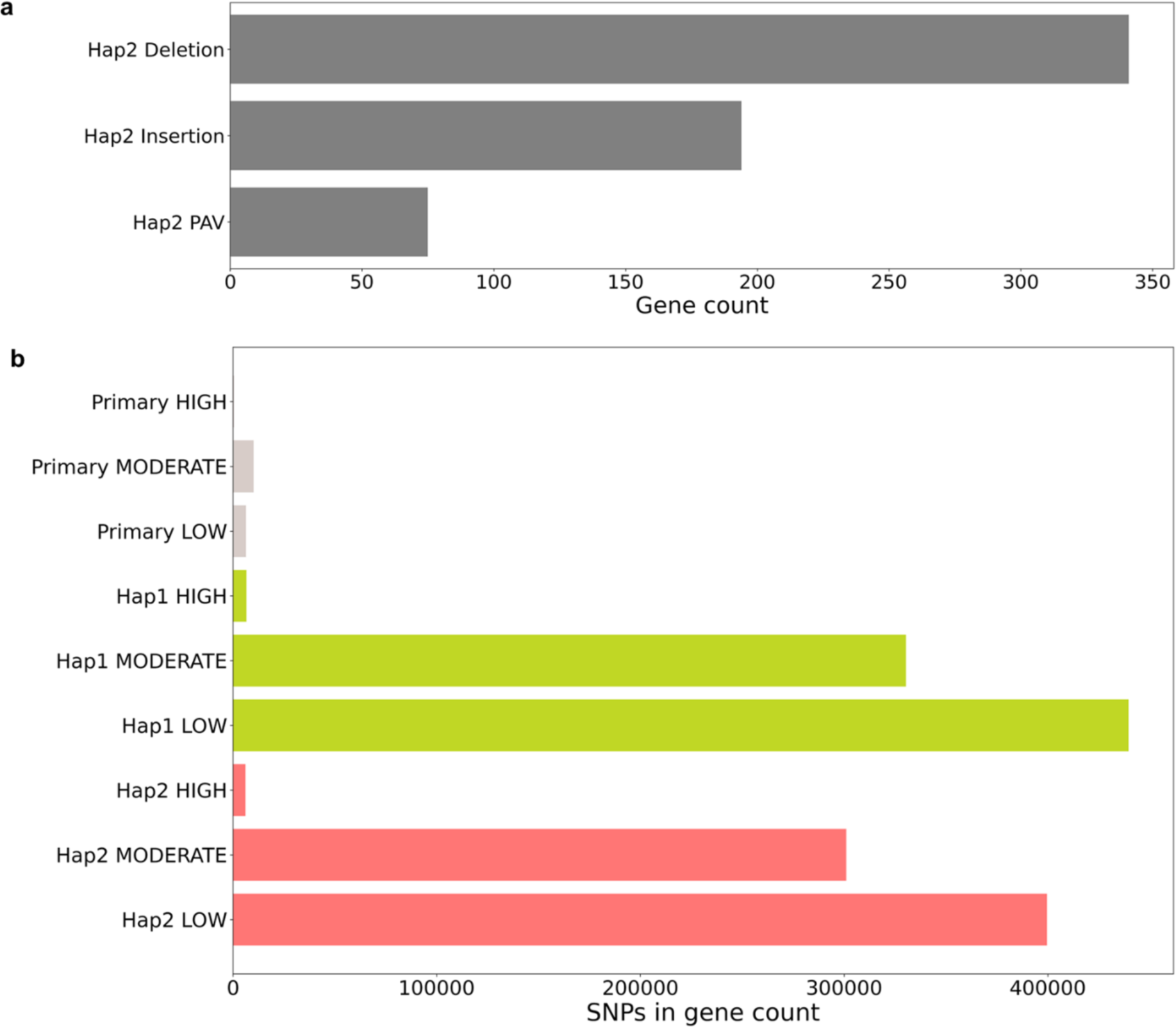
a) The number of genes related to SVs between Hap 1 and Hap 2. b) The number of SNPs related with genes in primary assembly, Hap 1, and Hap 2 genome.

### Identification of putative biosynthetic genes

The CDSs of all genes were functionally annotated based on similarity with universal protein knowledgebase (Bateman *et al*. 2017). The taxonomic descriptions of proteins were further utilized to select the candidate genes that are responsible for biosynthesis of roselle-enriched natural products: phenylpropanoids and flavonoids (Da-Costa-Rocha *et al*. 2014; Kim *et al*. 2021). The biosynthetic pathways of key intermediates involved in phenylpropanoids and flavonoids have been well-characterized (Dewick; Kanehisa and Goto 2000). Cinnamoyl-CoA and *p*-coumaroyl-CoA are the core frameworks of phenylpropanoids. Kaempferol and dihydrotricetin are known as the core intermediates of flavonoids. The candidate genes responsible for producing those intermediates were conserved in *H. sabdariffa* genome (Fig. 4 & Supplementary Data 2). To understand the biosynthetic pathway of roselle-enriched natural products, the first step would involve experimentally validating the catalytic activity of candidate genes. Furthermore, those genes could be used as baits in co-expression analysis to identify uncharacterized biosynthetic genes for roselle-enriched natural products such as hibiscetin (Fig. 4) (Da-Costa-Rocha *et al*. 2014; Kang *et al*. 2022).

**Figure 4.**
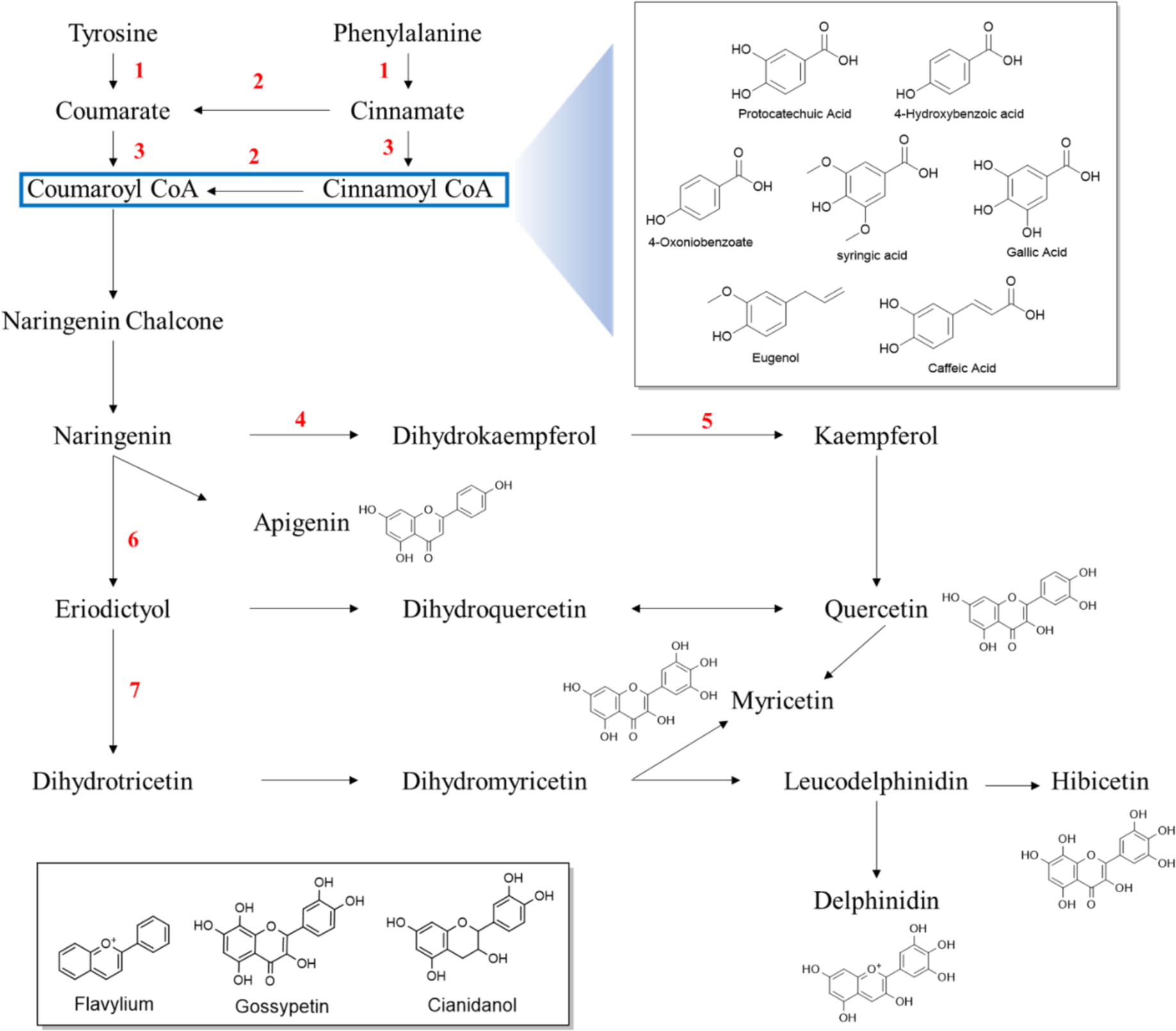
Phenylpropanoids and flavonoids of *H. sabdariffa*. The name and number of biosynthetic candidate genes in Hap 2 corresponding to each black arrow are following: 1, phenylalanine ammonia-lyase (14); 2, trans-cinnamate 4-monooxygenase (3); 3, 4-coumarate—CoA ligase (8); 4, 2-oxoglutarate 3-dioxygenase (4); 5, flavonol synthase (7); 6, flavonoid 3’-monooxygenase (8); 7, flavonoid 3’,5’-hydroxylase (8).

## Conclusion

Roselle is an important medicinal plant producing diverse natural products. However, the lack of genomic resource had limited the biological understanding of roselle and its pharmacological applications. In this study, we report the high-quality genome of *H. sabdariffa* and putative genes related with its secondary metabolite biosynthesis. Our results could be a foundation for understanding genetic evolution and metabolism in roselle.

